# Accurate and robust inference of microbial growth dynamics from metagenomic sequencing

**DOI:** 10.1101/2021.02.02.429365

**Authors:** Tyler A. Joseph, Philippe Chlenski, Tal Korem, Itsik Pe’er

## Abstract

Patterns of sequencing coverage along a bacterial genome—summarized by a peak-to-trough ratio (PTR)—have been shown to accurately reflect microbial growth rates, revealing a new facet of microbial dynamics and host-microbe interactions. Here, we introduce CoPTR (Compute PTR): a tool for computing PTRs from complete reference genomes and assemblies. We show that CoPTR is more accurate than the current state-of-the-art, while also providing more PTR estimates overall. We further develop theory formalizing a biological interpretation for PTRs. Using a reference database of 2935 species, we applied CoPTR to a case-control study of 1304 metagenomic samples from 106 individuals with irritable bowel disease. We show that PTRs have high inter-individual variation, are only loosely correlated with relative abundances, and are associated with disease status. We conclude by demonstrating how PTRs can be combined with relative abundances and metabolomics to investigate their effect on the microbiome.

**Availability:** CoPTR is available from https://github.com/tyjo/coptr, with documentation on https://coptr.readthedocs.io.

## 1 Introduction

Dynamic changes in the human microbiome play a fundamental role in our health. Understanding how and why these changes occur can help uncover mechanisms of disease. In line with this goal, the Integrative Human Microbiome Project and others have generated longitudinal datasets from disease cohorts where the microbiome has been observed to play a role [1–5]. Yet, investigating microbiome dynamics is challenging. On one hand, a promising line of investigation uses time-series or dynamical systems based models to investigate community dynamics [6–11]. On the other hand, the resolution of such methods is limited by sampling frequency, which often has physiological constraints on sample collection for DNA sequencing. Furthermore, while such methods accurately infer changes in abundance, they do not assess changes in growth rates.

Korem et al. [12] introduced a complementary approach to investigate microbiome dynamics. They demonstrated that sequencing coverage of a given species in a metagenomic sample reflects its growth rate. They summarized growth rates by a metric called the peak-to-trough ratio (PTR): the ratio of sequencing coverage near the replication origin to the replication terminus. Thus, PTRs provide a snapshot of population growth at the time of sampling, and their resolution is not limited by sampling frequency.

Their original method—PTRC—estimates PTRs using reads mapped to complete reference genomes. It has been used as a gold standard to evaluate other methods [13–15]. However, most species lack complete reference genomes, reducing its utility to researchers in the field. Therefore, follow-up work has focused on estimating PTRs from draft assemblies: short sections of contiguous sequences (contigs), where the order of contigs along the genome is unknown. Each approach relies on reordering binned read counts or contigs by estimating their distance to the replication origin. Although less accurate than PTRC, these methods allow PTRs to be estimated for a larger number of species. iRep [13] sorts binned read counts along a 5Kb sliding window, then fits a log-linear model to the sorted bins to estimate a PTR. GRiD [14] sorts the contigs themselves by sequencing coverage. It fits a curve to the log sequencing coverage of the sorted contigs using Tukey’s biweight function. DEMIC [15] also sorts contigs. However, it uses sequencing coverage across multiple samples to infer a contig’s distance from the replication origin. Specifically, DEMIC performs a principal component analysis on the contig by log2 coverage matrix across samples. The authors demonstrate that the scores along the first principal component correlate with distance from the replication origin. Ma et al. [16] provide theoretical criteria for when such an approach is optimal. Finally, other estimators have focused on PTR estimation for specific strains [17], or estimation using circular statistics [18].

Nonetheless, using PTRs has several limitations. From a theoretical perspective, it is not clear what PTRs estimate and how they should be interpreted. Bremer and Churchward [19] demonstrated that under exponential growth PTRs measure chromosome replication time and generation time, but this must be checked under arbitrary models of dynamics. From a practical perspective, estimating PTRs at scale requires running multiple tools across multiple computational environments—a cumbersome task.

In the present work we seek to address these issues. Our contributions are threefold. First, we provide theory that shows PTRs measure the rate of DNA synthesis and generation rate, regardless of the underlying dynamic model. Second, we derive two estimators for PTRs—one for complete reference genomes and one for draft assemblies. Third, we combined our estimators in a easy-to-use tool called CoPTR (Compute PTR). CoPTR provides extensive documentation, a tutorial, and a precomputed reference databases for its users. We demonstrate that CoPTR is more accurate than KoremPTR—a reimplementation of PTRC—on complete reference genomes, and more accurate than the current state-of-the-art on metagenome-assembled genomes (MAGs). We conclude with a large scale application to a dataset of 1304 metagenomic samples from a case-control cohort of individuals with irritable bowel disease [3].

## 2 Results

### 2.1 CoPTR Overview

The method we developed models the density of reads along the genome in a sample by adapting an argument proposed by Bremer and Churchward [19]. Under an assumption of exponential growth, they showed that the copy number ratio of replication origins to replication termini in a population, *R*, is given by

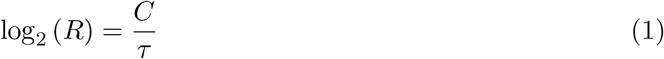

where *C* is the time it takes to replicate a bacterial chromosome, and *τ* is the (fixed) generation time. We generalize this (see Supplementary Note 1) for dynamic quantities:

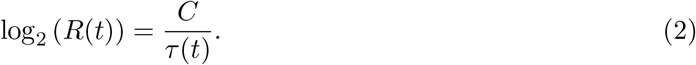

The variable *τ* now depends on collection time *t*. When a complete reference genome is available the PTR is an estimator for *R*(*t*). However, the PTR is only correlated with *R*(*t*) on draft assemblies because the assembly may not include the replication origin or terminus.

The derivation also suggests that copy number along the chromosome decays log-linearly away from the replication origin (Supplementary Note 2). We used this fact to develop CoPTR (Compute PTR): a maximum likelihood method for estimating PTRs from complete genomes and draft assemblies (Figure 1). CoPTR takes sequencing reads from multiple metagenomic samples and a reference database of complete and draft genomes as input. It outputs a genome by sample matrix where each entry is the estimated log_2_(PTR) for each species in that sample. It has two modules: CoPTR-Ref that estimates PTRs from complete genomes, and CoPTR-Contig that estimates PTRs from draft assemblies. As such, it combines the improved accuracy enabled by complete genomes with the flexibility afforded by being able to work against draft and metagenomic assemblies. For both methods, sequencing reads are first mapped to the reference database. CoPTR-Ref estimates PTRs by applying an adaptive filter to remove regions of ultra-high or ultra-low coverage. Then it fits a probabilistic model to estimate the PTR and replication origin. CoPTR-Contig estimates PTRs by first binning reads into approximately 500 non-overlapping windows. It filters out windows with excess or poor numbers of reads. Coverage patterns across multiple samples are used to reorder bins using Poisson PCA. The reordered bins serve as approximate genomic coordinates which are used to obtain maximum likelihood estimates of PTRs.

**Figure 1:**
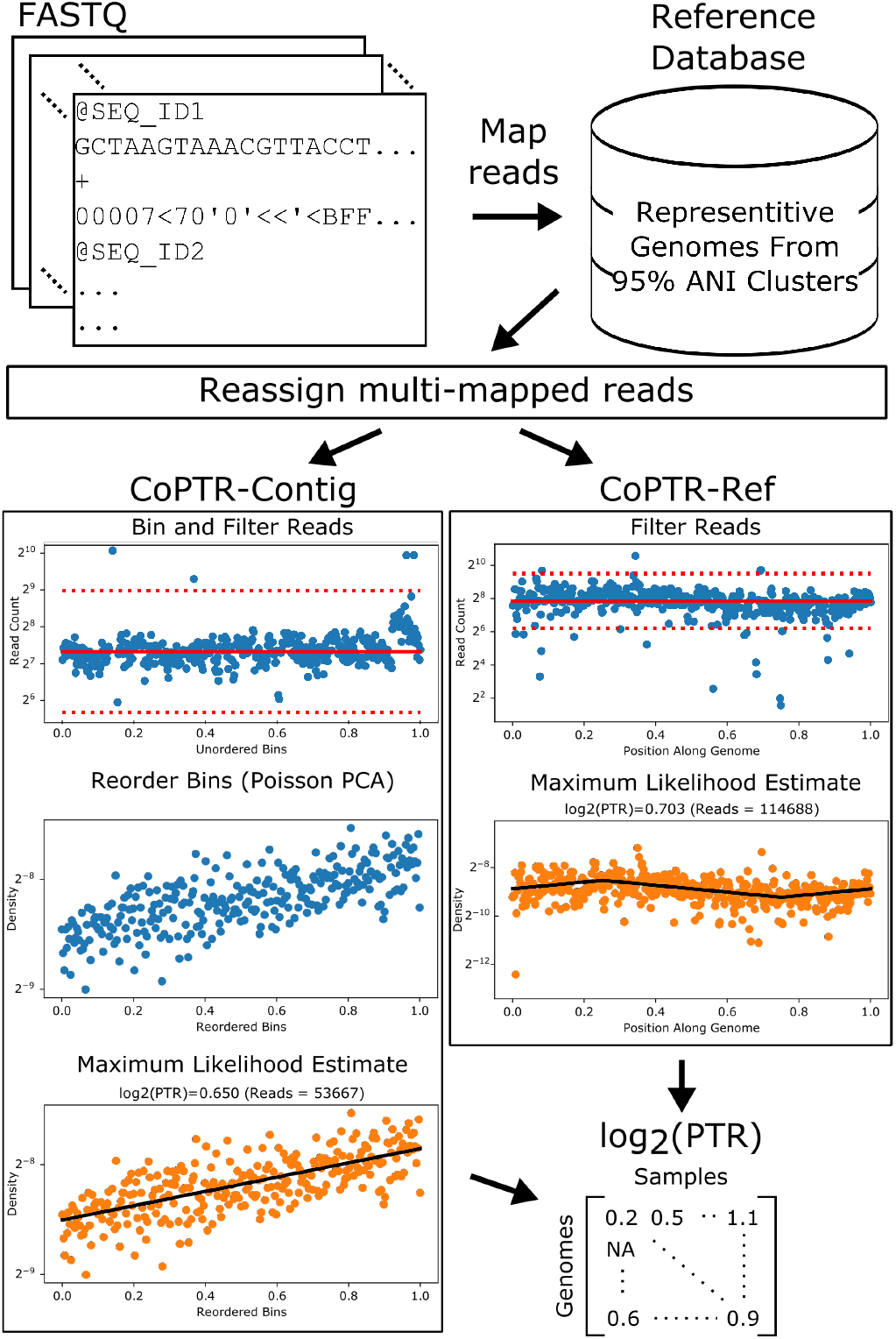
CoPTR Workflow. Sequencing reads from multiple metagenomic samples are mapped to a reference database containing representative strains from complete reference genomes and high-quality assemblies (*>* 90% completeness, *<* 5% contamination). Multi-mapped reads are reassigned to a single genome using a probabilistic model. Then, PTRs are computed for each genome in each sample. For species with complete reference genomes, PTRs are estimated by maximizing the likelihood of a model describing the density of reads along the genome (CoPTR-Ref). For species with high-quality assemblies, reads are binned across the assembly, bins are reordered based on sequencing coverage across multiple samples using Poisson PCA, and the slope along this order is estimated by maximum likelihood (CoPTR-Contig). CoPTR outputs a table of the log_2_(PTR) per genome in each sample.

### 2.2 CoPTR-Ref accurately estimates PTRs using complete reference genomes

We evaluated CoPTR-Ref on simulated data. Briefly, we simulated read counts based on read density maps generated from high coverage genomic samples of *Escherichia coli, Lactobacillus gasseri*, and *Enterococcus faecalis* from Korem et al. [12] (Supplementary Figure 1). The density maps reflect differences in coverage along a genome due to GC content and mappability. To facilitate comparison with CoPTR-Ref, we also reimplemented PTRC using code provided by the authors. The new implementation, called KoremPTR, was designed to work with simulated read counts and reads mapped with Bowtie2. KoremPTR showed a good correspondence with the original method (Pearson *r >* 0.99; Supplementary Figure 2).

Notably, our simulations demonstrated that CoPTR-Ref more accurately estimates PTRs than KoremPTR (Figure 2, Supplementary Figure 3), requiring as few as 5000 reads to achieve greater than 0.95 Pearson correlation across density maps (Supplementary Figure 3). KoremPTR appeared to underestimate the simulated PTR, causing the difference in accuracy (Figure 2B, Supplementary Figure 3). Nonetheless, PTR estimates by KoremPTR were highly correlated with the ground truth (Pearson *r >* 0.88). We saw the same pattern across 6 genomic (bacteria grown in monoculture) and metagenomic datasets (Figure 2C). Both methods were correlated, but CoPTR-Ref estimated larger PTRs than KoremPTR on the same samples.

**Figure 2:**
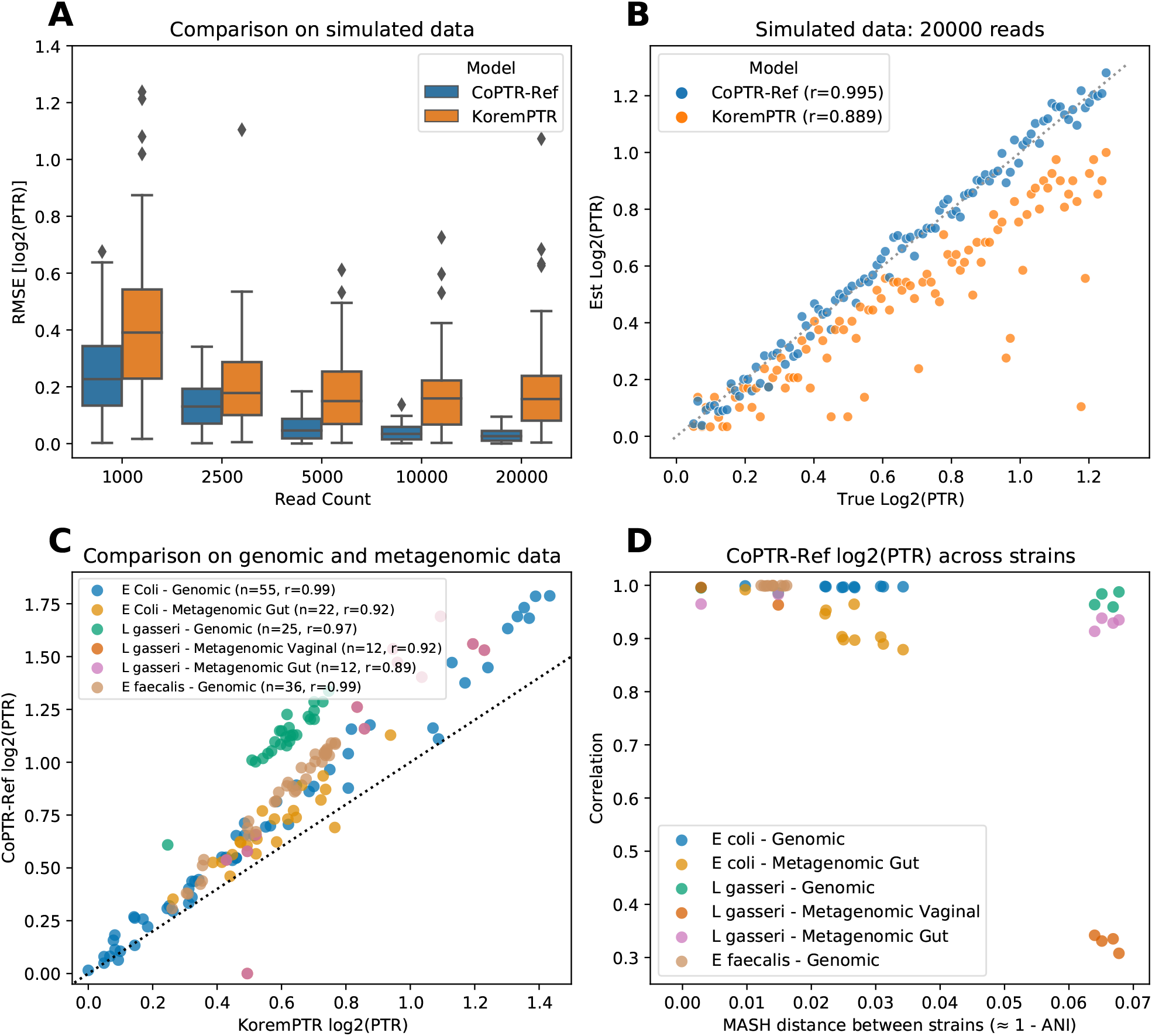
CoPTR-Ref is accurate on simulated and real data. **(A)** Accuracy of CoPTR-Ref and KoremPTR on simulated data based on an *E. coli* genome. Performance was compared by computing the root-mean-square-error (RMSE) of the log_2_(PTR) (*y*-axis) across 100 replicates while varying the number of reads (*x*-axis), varying the position of the replication origin, and varying the PTR. **(B)** Ground truth (*x*-axis) and estimated (*y*-axis) log_2_(PTR) across 100 simulation replicates with 20000 reads. KoremPTR appears to underestimate the true log_2_(PTR). **(C)** Comparison of KoremPTR log_2_(PTR) (*x*-axis) and CoPTR log_2_(PTR) on 6 real genomic and metagenomic datasets. **(D)** Evaluation of CoPTR-Ref’s log_2_(PTR) estimates using representative genomes from different strains (5 *E. coli* strains, 4 *L. gasseri* strains, and 5 *E. faecalis* strains). Each dataset in panel C was mapped to strains from the same species, and the Pearson correlation (*y*-axis) was computed for each pair of strains. When the distance between strains (*x*-axis) is small, log_2_(PTR)’s are highly correlated.

To evaluate whether variation among representative genomes per 95 % ANI clusters—an operational threshold for defining species [20]—affects the accuracy of CoPTR, we mapped the same samples to different strains. We found that PTR estimates were robust to strain variation when the MASH distance [21] between strains was less than 0.05—corresponding to ∼ 95% ANI (Figure 2D). These results indicate that one reference genome per 95% ANI cluster can be included in a reference database without loss of information.

We also compared log_2_(PTR) estimates to changes in population size of *E. coli* grown in culture. If *N* (*t*) is the size of the population at time *t*, our theory suggests that in this restricted setting 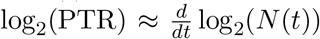. We found a strong correlation (*r >* 0.96) between log_2_(PTR) and a finite difference estimate of log_2_(*N* (*t*)) computed from optical density measurements of the culture (Supplementary Figure 4).

### 2.3 CoPTR-Contig accurately estimates PTRs using MAGs

Because CoPTR-Contig reorders bins, not contigs, we could directly compare CoPTR-Ref to CoPTR-Contig using the same simulation framework (Figure 3A, Supplementary Figure 5). Estimates by CoPTR-Contig were highly correlated (Pearson *r >* 0.9) with the simulated ground truth with as a few as 5000 reads, but were overall less accurate than CoPTR-Ref. Our results highlight the benefit of using the additional information provided by complete reference genomes.

**Figure 3:**
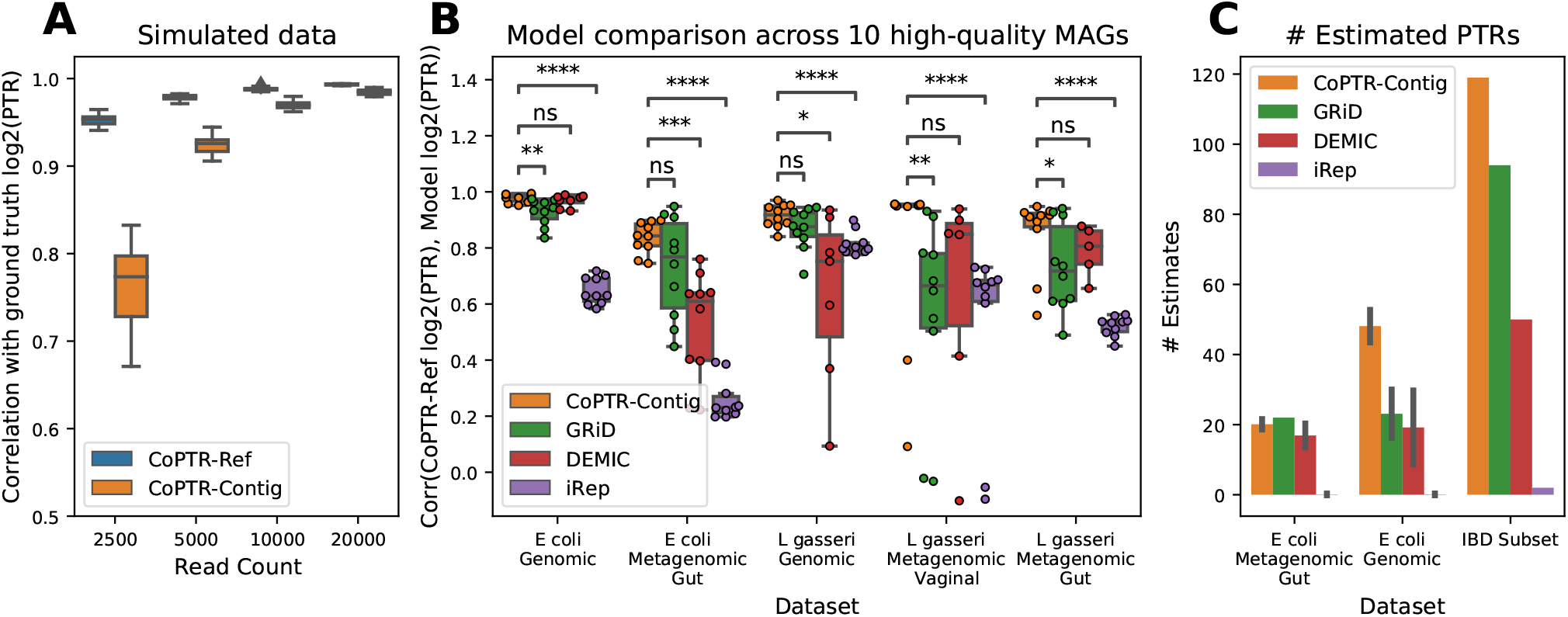
CoPTR-Contig is accurate on simulated and real data. **(A)** Comparison of CoPTR-Ref and CoPTR-Contig on simulated data using the *E. coli* density map. Performance was evaluated by computing the correlation (*y*-axis) between simulated and estimate log_2_(PTR)’s across read counts (*x*-axis), randomly chosen replication origins, and PTRs. CoPTR-Contig shows high accuracy above 5000 reads. **(B)** Comparison of CoPTR-Contig to GRiD, DEMIC, and iRep across 5 genomic (monoculture) and metagenomic datasets (*x*-axis). For each dataset, reads were mapped to a single reference genome for each species (see Methods). Performance was evaluated by comparing log_2_(PTR) estimates from CoPTR-Ref to the log_2_(PTR) estimate from each method across 10 high-quality metagenome assembled genomes (MAGs; points on the figure). Significance was computed using a two-tailed *t*-test (*: *p <* 0.05; **: *p <* 10^*−*2^; ***:*p <* 10^*−*3^; ****: *p <* 10^*−*4^). **(C)** Number of PTR estimates from species passing the filtering criteria for each model. The mean and standard deviation are reported for the *E. coli* metagenomic gut and genomic datasets across MAGs from (B). Error bars depict one standard deviation. Each model was also applied to 10 samples from the IBD dataset (Section 1) using 1,009 high-quality MAGs from the IGGdb. The total number of PTRs passing filtering criteria for each model is reported.

To assess the applicability of our method to metagenomic assemblies, which are of variable quality and contamination levels, we performed simulations investigating the impact on the accuracy of CoPTR-Contig. We found that CoPTR-Contig is robust to the level of genome completeness, providing comparable accuracy with completeness as low as 50%. We further found that CoPTR-Contig’s estimates are robust to moderate amounts of up to 5% contamination in the assembly from other species (Supplementary Figure 6).

We then compared CoPTR-Contig to GRiD, DEMIC, and iRep across 5 real genomic and metagenomic datasets of *E. coli* and *L. gasseri* where both complete reference genomes and metagenomic assembled genomes (MAGs) were available (Figure 3B). We considered 10 high-quality MAGs (*>* 90% completeness *<* 5% contamination) from the IGGdb [22] and computed the correlation between the log_2_(PTR) estimate from each method and the log_2_(PTR) from CoPTR-Ref. For CoPTR-Ref, reads were mapped to a single complete genome (see Methods). All 10 of the *E. coli* MAGs were assigned to the same 95% ANI species cluster, while 8 of the 10 *L. gasseri* MAGs were from one cluster, and the remaining 2 from another. To allow for a fair comparison, we changed the default parameters of each method to allow estimates on each sample—with the exception of DEMIC which provides no command line options to change filtering criteria. We note that almost all the samples we explored were below the minimum recommended coverage for iRep (Figure 3C, Supplementary Figure 7).

We found that CoPTR-Contig significantly outperformed (*p*-value *<* 0.05 using a two-sided paired *t*-test; the 2 *L. gasseri* MAGs from a different species cluster were excluded) GRiD, DEMIC, and iRep on 3, 2, and 5 of the datasets respectively. All models performed poorly on the 2 *L. gasseri* MAGs that were from a different 95% ANI cluster (outliers on Figure 3B), recapitulating results from the strain comparison experiment using CoPTR-Ref (Figure 2D). Many of the comparisons between CoPTR-Contig and DEMIC failed to reach significance because DEMIC estimated fewer PTRs overall (Figure 3C, Supplementary Figure 7), resulting in fewer MAGs for comparison (points in Figure 3C).

An important aspect affecting the utility of PTR inference methods is the number of PTR estimates they are able to provide for a given sample. We therefore compared the number of estimated PTRs that passed the filtering criteria of each method (Figure 3C, Supplementary Figure 7). We mapped 10 samples from the IBD dataset (Section 2.5) to 1,009 high-quality MAGs from the IGGdb, and counted the number of PTR estimates. The reported estimates for GRiD are based on GRiD’s published minimum coverage requirement: species with *>* 0.2x sequencing coverage. We were unable to run GRiD’s high-throughput model on two systems (Ubuntu 18.04.4 LTS and macOS 10.15) to produce estimates on this dataset. We found that CoPTR-Contig produced more PTR estimates overall than the other models we evaluated. Importantly, this number does not include the additional estimates from complete genomes using CoPTR-Ref. Taken together with the improved accuracy of CoPTR (Figure 3B), these results show that CoPTR outcompetes previous PTR estimation methods in both the number of estimates produced and their accuracy, demonstrating its utility for microbiome analysis.

### 2.4 PTRs recapitulate a signal of antibiotic resistance

We next evaluated if we could use CoPTR to detect a signal of antibiotic resistance in *Citrobacter rodentium*. Korem et al. [12] generated 86 samples from 3 populations of *in vitro* culture of *C. rodentium*. One population was treated with Erythromycin, a growth inhibiting antibiotic; another was treated with Nalidixic acid to which *C. rodentium* is resistant. The final population was a control and received no treatment.

We wanted to see if we could recapitulate this signal using CoPTR. Similar to the original study, we observed a difference in PTRs between the populations exposed to Erythromycin and Nalidixic acid (Supplementary Table 1). In addition, our results add to the original study by assigning an effect size to each condition. We found that Erythomycin has a strong negative effect size on the log_2_(PTR), while Nalidixic acid has a strong positive effect size.

### 2.5 PTRs are highly personalized

We next sought to demonstrate how PTR measurements can be used in a large-scale study. To this end, we considered 1304 metagenomic samples from 106 individuals in a case-control study of irritable bowel disease (IBD) [3]. Individuals in the study had two different subtypes of IBD: Crohn’s disease and Ulcerative colitis. We mapped the metagenomic samples to a database from IGGdb [22] consisting of 2935 complete genomes, assemblies, and MAGs, selected as representative genomes from 95 % ANI clusters.

A large dataset with multiple samples per individual allowed us to investigate questions about sources of variation for PTRs. Thus, we estimated the fraction of variation explained by differences between individuals, disease-statuses, ages, and sex. Inter-individual differences in PTRs accounted for the largest fraction of variance largest among variables explored (Figure 4A), consistent with the original study that found inter-individual variation was the largest source of variation among the other multi-omic measure types collected [3]. Notably, PTRs were mostly uncorrelated with relative abundances, suggesting that PTRs tag a signal of biological variation complementary to relative abundances (Figure 4B).

**Figure 4:**
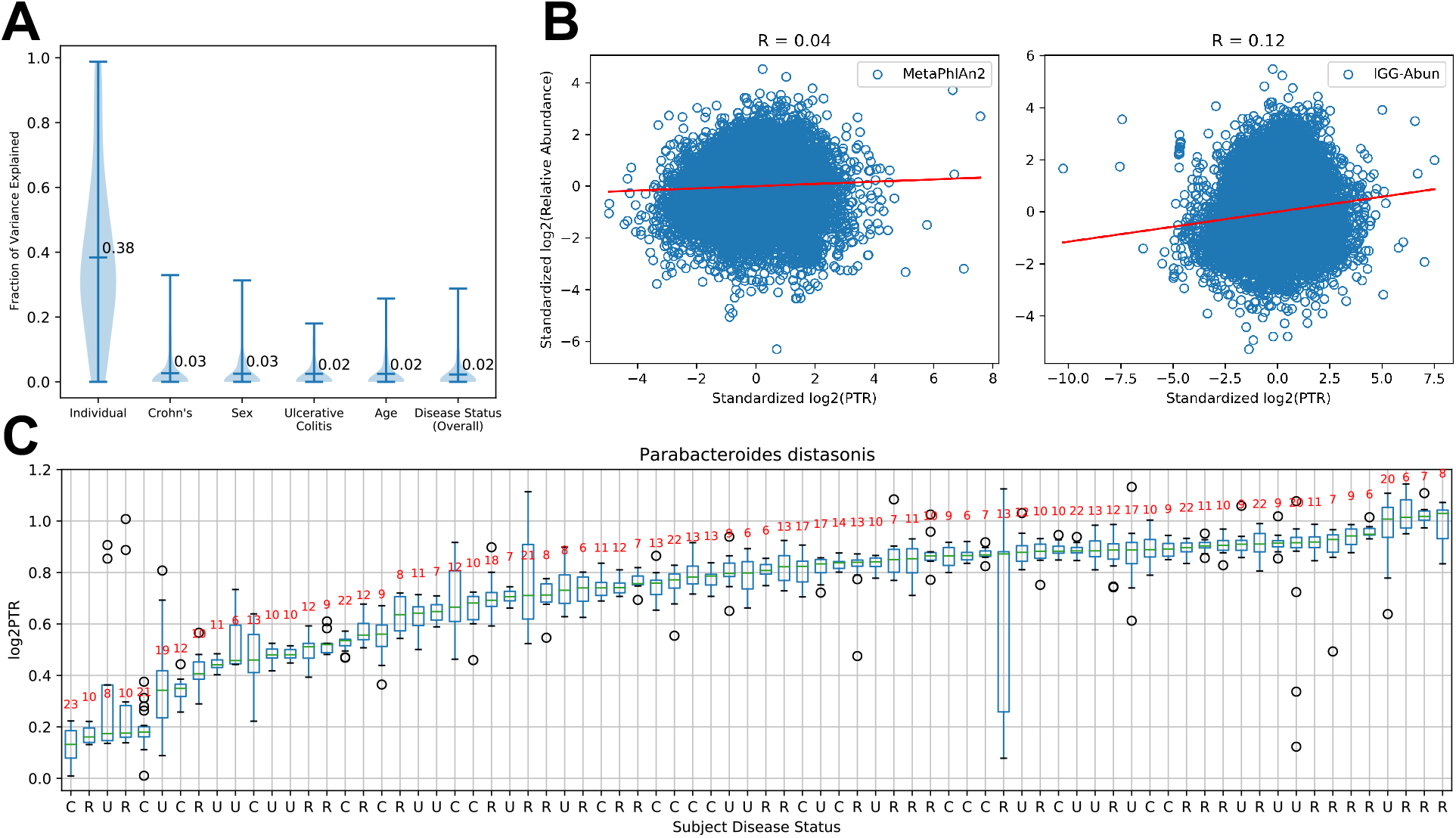
PTRs are highly personalized and uncorrelated with relative abundances. **(A)** Fraction of variance of log_2_(PTR) explained per species by variation between individuals, disease-statuses, age, and sex. Inter-individual variation accounts for most variation in among log_2_(PTR)s of the variables explored. **(B)** Correlation between standardized log_2_(PTR) and log_2_(relative abundance) on species matched to relative abundances from MetaPhlAn2 (left) and directly estimated from read counts from reads mapped to the IGGdb reference database (right). **(C)** Boxplots of the log 2(PTR) (*y*-axis) of *Parabacteroides distasonis* across individuals (*x*-axis). *P. distasonis* was the most significant species when testing for individual differences using the Kruskal-Wallis test on controls only. Individuals are labeled by disease status (C: control; R: Crohn’s disease; U: Ulcerative colitis), and the *y*-axis are the log_2_(PTR) per individual. Sample sizes are displayed in red. In most individuals the PTR of *P. distasonis* exhibits only small variations among samples, but large variation across individuals.

### 2.6 PTRs are associated with IBD

We then asked if we could associate species to disease status through their PTRs. We found 1 species that was significantly associated (FDR *q* = 0.025, effect size = −0.1574) with Crohn’s disease (Supplementary Table 2), *Subdoligranulum sp*., and three species with Ulcerative colitis (Supplementary Table 3): *Roseburia intestinalis* (*q* = 1.07 × 10^*−*3^, effect size = 0.094), *Ruminiclostridium sp* (*q* = 2.5 × 10^*−*2^, effect size = −0.138), and *Subdoligranum sp*. (*q* = 2.69 × 10^*−*2^, effect size = −0.168). (Figure 5A). Notably, Vila et al. [23] also report an increased PTR in *R. intestinalis* in individuals with Crohn’s disease and Ulcerative colitis in a separate cohort, using PTRC. We did not not observe a significant association between the relative abundance of *R. intestinalis* and disease status, nor did Vila et al. [23]. Altogether, our results provide additional evidence that *R. intestinalis* may play a role in Ulcerative colitis, observable only through analysis of growth dynamics.

For the remaining investigation we focused on *R. intestinalis*. We asked if we could assess the impact of various species on *R. intestinalis* by associating relative abundances across species estimated with MetaPhlAn2 [24] with its log_2_(PTR) (Supplementary Table 4). We found two species with a positive association with *R. intestinalis*, and one with a strong negative association (Figure 5B). Finally, we investigated if we could relate metabolomic measurements to log_2_(PTR)s (Supplementary Table 5). We found two metabolites with a positive association with the log_2_(PTR) of *R. intestinalis*. Notably, one of them—2-hydroxyglutarate—is part of the butanoate metabolic pathway, and *R. intestinalis* is a known butyrate producing bacteria.

**Figure 5:**
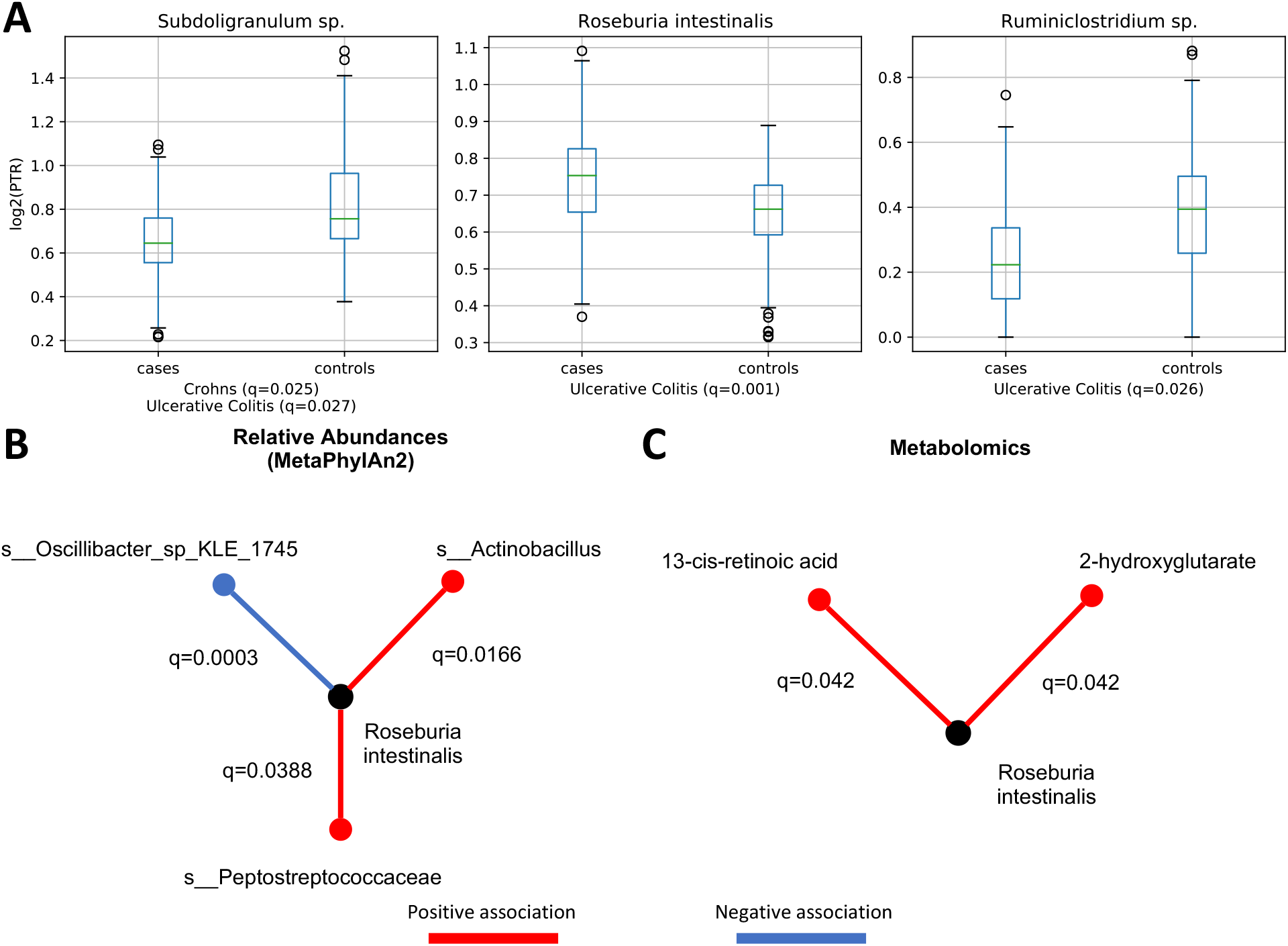
Association of log_2_(PTR)s with disease status (A), relative abundances (B), and metabolomics (C). **(A)** log_2_(PTR)*s* can be used to associate species with disease status. Significance was assessed by a fitting a linear model to log_2_(PTR) per species and correcting for false-discoveries (*q*-values denote false-discovery rate). PTRs can be combined with relative abundances to assess species interactions **(B)** or the impact of metabolites **(C)**.

## 3 Discussion

Peak-to-trough ratios (PTRs) have the potential to be a valuable tool for investigating microbiome dynamics. Here, we provided theory giving PTRs a biological interpretation. We introduced CoPTR, a software system combining two methods for estimating PTRs: CoPTR-Ref estimates PTRs with the assistance of a complete reference genome, and CoPTR-Contig estimates PTRs from draft assemblies. We showed that CoPTR-Ref is more accurate than KoremPTR, the current gold standard for PTR estimation from complete reference genomes. We also showed that CoPTR-Contig was more accurate than the current state-of-the-art for PTR estimation using draft assemblies, while providing more PTR estimates overall. Importantly, CoPTR is easy to use, has extensive documentation, and provides a precomputed reference database for its users.

When building CoPTR we focused on estimating PTRs per species, rather than per strain. Our goal was to allow CoPTR to be applied to recent database efforts that combined representative genomes from MAGs, assemblies, and complete genomes clustered at approximately 95% average nucleotide identity [25–27, 22, 28, 29]. There are benefits and drawbacks to this approach. The major benefit is reduction in database size, and therefore in computational time required for read mapping. The larger IGGdb database of all high-quality gut MAGs from Nayfach et al. [22] contains 24,345 genomes which is considerable larger than the 2935 genomes used here. Our results showed that PTR estimates from the same samples mapped to different closely related strains were highly concordant. Thus, there is not much to be gained from including all strains in the reference database. Nonetheless, the drawback is that CoPTR may not distinguish differences in PTRs across samples due to differences in strains.

We also focused on estimating PTRs from high-quality MAGs (*>*90% completeness, *<*5% contamination). Inference from MAGs is more challenging than other assembly types, due to differences in assembly completeness and contamination from other species. Many things can go wrong during the assembly processes. These, in turn, can affect PTR inference. In our opinion, it is better to have fewer high-quality estimates than more poor-quality ones, and for this reason we have chosen strict inclusion criteria for MAGs.

Our results on the IBD dataset showed that PTRs were highly personalized, mirroring results from other measurements in the original study. The largest fraction of variance observed was attributable to inter-individual variation. Additionally, PTRs were uncorrelated with relative abundances. These facts combined suggest that PTRs are tagging some source of biological variation not captured by other measurement types, and can complement other approaches for interrogating the microbiome.

There are other benefits to using PTRs as well. Compared to relative abundances, PTRs have a clearer biological interpretation, because an increase in relative abundance does not necessarily correspond to an increase in population size. In contrast, we showed that an increase in PTR in a species corresponds to an increase in the rate of DNA synthesis, and that an increase in the log PTR corresponds to a decrease in generation time. Either of these facts can be used to generate hypotheses about the drivers of differences across conditions. Furthermore, because PTRs provide a snapshot of growth at the time of sampling, they potentially alleviate the need to perform dense-in-time sampling typically needed to detect dynamic changes. This suggests that it may be more cost-effective to sequence more individuals, rather than more samples per individual. Finally, we showed that relative abundances and metabolomic profiles can be used to associate species or metabolites with PTRs. Altogether, our study demonstrates that PTRs can provide new approaches for investigating community interactions, relating multiomic measurements to the microbiome, and for investigating the relationship between microbiome dynamics and disease.

## 4 Methods

### 4.1 CoPTR Implementation

#### Read mapping

Reads are mapped using Bowtie2 [30] using the parameter `-k 10` to allow up to 10 mappings per read. We chose this parameter after observing that 99% of reads mapped to 10 or fewer locations on the IGGdb using a subset of 10 samples from the IBD dataset. Reads with fewer than 10 mapping were assigned using a variational inference algorithm described in Supplementary Note 3. In present work, reads with 10 (or more) mappings were discarded from downstream analysis. However, CoPTR has a command line argument to adjust this setting.

Before reassigning multiply mapped reads, reads are filtered by alignment score. Alignment score is more sensitive than mapping quality, since different alignment scores can result in the same mapping quality. Bowtie2 assigns penalties to mismatched bases weighted on their quality score. Bases with a perfect quality score receive a -6 penalty for a mismatch, decreasing as the quality score decreases. For a read of length *L*, we filtered out reads with a score less than −6\**L*\*(1−0.95). Given a read with perfect quality scores, this corresponding to removing reads with less than 95% identity to the reference sequence. Of course reads do not have perfect quality scores, so this threshold is less strict than 95% identity.

#### CoPTR-Ref

PTRs from species with complete reference genomes are estimated with CoPTR-Ref. Regions of the genome with excess or poor coverage per sample are first filtered out in two steps. In the first step we apply a coarse-grained filter by binning reads into 500 bins. Let *m* be the median log_2_ read count across nonzero bins, and *s* the larger of 1 or the standard deviation of log_2_ read counts in nonzero bins. Bins are filtered out if they fall outside the interval (*m* − *α*_0.025_, *m* + *α*_0.025_), where *α*_0.025_ is the two-sided (1 − 0.025) critical region from an *N* (*m, s*) distribution. After the coarse-grained filter, we apply a fine-grained filter by computing read counts across a rolling window encompassing 12.5% of the genome. We apply the same filtering criteria around the center of each window.

After filtering, the remaining bins are concatenated, and read positions normalized so that they fall in the unit interval [0, 1]. Let *x ∈* [0, 1] be the coordinate of a read, *x*_*i*_ be the coordinate of the replication origin, and *x*_*t*_ = (*x*_*i*_ + 0.5) mod 1 be the replication terminus. We estimate the log_2_(PTR) and replication origin across all samples by maximizing the likelihood of the model

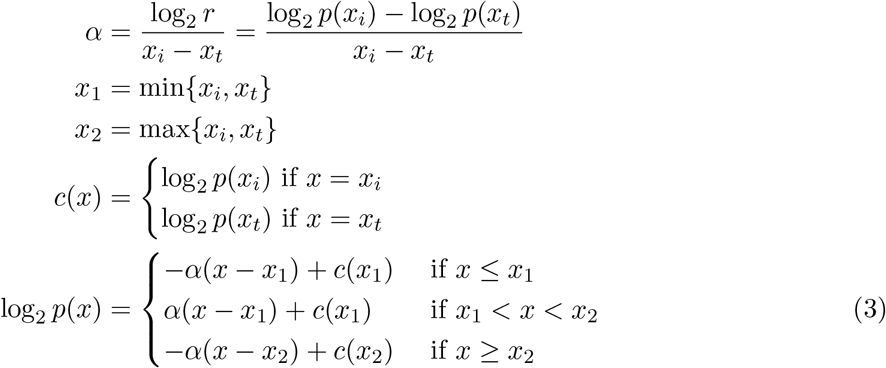

We describe how to compute log_2_ *p*(*x*_*i*_), log_2_ *p*(*x*_*t*_), and the normalizing constant in Supplementary Note 2. We maximize the likelihood using the `SLSQP` optimizer in `SciPy` [31]. We first maximize with respect to each sample separately to get initial estimates of the log_2_(PTR) per sample, then jointly estimate the replication origin given these estimates. Finally, given the estimated replication origin from all samples, each individual log_2_(PTR) is updated once more.

#### CoPTR-Contig

PTRs from species with draft assemblies are estimated with CoPTR-Contig. Reads across contigs are binned into approximately 500 bins (adjusted such that the average length of each bin is divisible by 100bp). We choose 500 bins, rather than fixed bin size, so that the model would behave similarly across genomes of different lengths. We then apply a similar coarse-grained filter to the log_2_ read counts binned into 500 bins. Bins that are filtered are marked as missing for the Poisson PCA step.

The remaining bins are reordered by applying a Poisson PCA to read counts across samples. Let *B* be the number of bins, and *N* the number samples. Let *x*_*bi*_ be the read count in bin *b* from sample *i*, and let *Ω* = {*x*_*bi*_ : bin *b* is not missing from sample *i*}. In Poisson PCA, we model the read count in each bin using a matrix *C* = *V W* ∈ ℝ^*B×k*^ × ℝ^*k×N*^ with low-rank structure. Specifically, we assume rank 1 structure where *W* ∈ ℝ^*B×*1^ and *V* ∈ ℝ^1*×N*^. The read count *x*_*bi*_ is modeled by

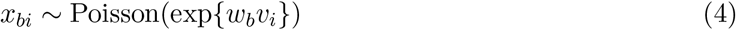

The parameters *W* and *V* are estimated by iteratively maximizing the likelihood

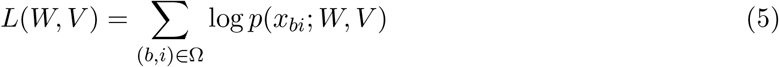

with respect to *W* then *V* until convergence.

The scores for each bin *w*_*b*_ are used to rank bins from low to high, representing approximate distance from the replication origin. Bins are reordered by their rank, then for each sample the top and bottom 5% of bins removed. The log_2_(PTR) is estimated by maximizing a discretized version of equation 3 using the `SLSQP` optimizing in `SciPy`, fixing the replication origin at one end and terminus at the other.

### 4.2 Simulations

To generate realistic simulations we computed read density maps by mapping reads from genomic (monoculture) samples to reference genomes where the strain was known. For each density map, we computed the read count in 100bp bins, then divided by the total number of reads to obtain empirical probabilities that a read originates from a location in the genome. These probabilities are conditioned on the PTR in the sample. We therefore used KoremPTR to estimate the PTR for each sample using the replication origin from the DoriC database [32], and reweighted the probabilities by the estimated PTR. Specifically, let *p*_1_, …, *p*_*N*_ be the unadjusted probabilities that a read originates from a bin, let 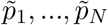 be the probabilities under the model given the replication origin and PTR, and let 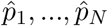 be the adjusted probabilities. The adjusted probabilities are

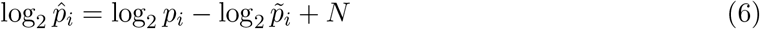

where *N* is the normalizing constant.

We generated density maps for *E. coli* from a genomic sample with 894,685 reads (14x coverage), a *L. gasseri* sample with 2,645,206 reads (104x coverage), and *E. faecalis* with 581,836 reads (14.75x coverage) from Korem et al. [12]. Supplementary Figure 1 displays the adjusted density maps. When simulating data, we performed the reversed adjustment by the simulated replication origin and PTR. Given 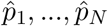, and theoretical probabilities for the simulated PTR and replication origin 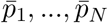, we computed the probability that a read is derived from bin *i* by computing

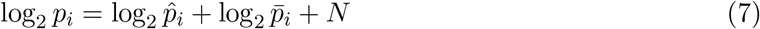

To compare CoPTR-Ref and KoremPTR, we performed 100 simulations each for read counts of 1000, 2500, 5000, 10000, and 20000. For each simulation, a random replication origin and PTR is chosen. Reads counts in 100 bp bins are simulated based on the adjusted probabilities described above, then converted to genomics coordinates. The coordinates are provided to CoPTR-Ref and KoremPTR to estimate PTRs.

To evaluate CoPTR-Contig, we performed 20 simulation replicates consisting of 100 samples each, while varying the number of simulated reads. Because PTR estimates can be sparse, we processed samples in batches of 5 to explore how well CoPTR-Contig reordered bins at small sample sizes.

#### Completeness and contamination experiments

We extended our simulation framework to investigate genome completeness and contamination using the *E. coli* density map to perform our simulations. To simulate genome completeness, we held our random fragments of the reference genome in 1% increments selected uniformly at random. The remaining sections of the genome were treated as contigs, and reads were simulated from the contigs. To simulate genome contamination, we simulated reads from two separate genomes: *E. coli* and *L. gasseri*. For a given contamination percentage *c*, reads were simulated from the *E. coli* genome, setting the completeness percentage to 100 − *c*. Then, simulated read counts from contigs in *L. gasseri* genome were added until the percentage of contamination by *L. gasseri* was *c*.

### 4.3 Datasets and reference genomes

We downloaded genomic samples from Korem et al. [12], and metagenomic samples from the Human Microbiome Project [33] and the IBD dataset [3]. Vaginal and gut metagenomic samples from the Human Microbiome Project were selected by mapping reads to reference genomes of *E. coli* and *L. gasseri*, and retaining samples with more than 2500 mapped reads. Gut samples of *L gasseri* from the IBD dataset were selected based on whether CoPTR had an estimated PTR. Complete accession numbers per experiment are listed in Supplementary Table 6.

To compare estimates across reference genomes, we downloaded reference genomes from NCBI. Accession numbers for genomes and MAGs are listed in Supplementary Table 6. We selected genomes from each of *E. coli, L. gasseri*, and *E. faecalis* matching the strains reported by Korem et al. [12], and performed comparison on genomic samples using these strains. Distances between reference genomes were computed using MASH [21]. The genomes NC 007779.1, NC 008530.1, and NZ CP008816.1 corresponds to the strains used by Korem et al. [12].

To compare estimates across MAGs, we downloaded high-quality assemblies from Nayfach et al. [22]. On both complete references and MAGs, we noted for *L. gasseri* that genomes were from two different 95% ANI species clusters. For the MAGs, 8 were from one cluster and 2 from another. To compare PTR estimates from *L. gasseri* MAGs to CoPTR-Ref estimates, we selected a reference genome corresponding to the species cluster with 8 MAGs. We did this by downloading a complete genome in the same species cluster identified by Nayfach et al. [22], and computing the MASH distance with genomes above. We found one genome with 0 MASH distance to the species cluster which we used for analysis.

When performing the model comparison and the *C. rodentium* experiments, we mapped reads to one genome at a time using Bowtie2’s default parameters.

### 4.4 Antibiotic resistance experiment

We applied CoPTR to a dataset of 86 longitudinal samples from three populations of *C. rodentium*. Samples were taken from three periods of the experiment: a treatment period where the antibiotic was applied, a recovery period when the antibiotic was removed, and a stationary period. The structure of the experiment requires variables to account for the sampling time under each period. Let 𝒫= {Treatment, Recovery, Stationary}, and for each *p* ∈ 𝒫 denote *T*_*p*_ as the number of time points. We fit the following model:

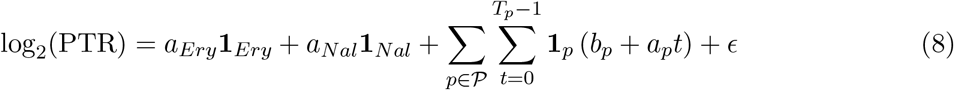

The parameters *b*_*p*_ say that the mean log_2_(PTR) differs under each period, and *a*_*p*_ model directional changes within the period over time. The variables *a*_*Ery*_ and *a*_*Nal*_ measure the effect of each antibiotic on the log_2_(PTR). While the model is somewhat complex, it is a reflection of the sampling process and dynamics of *in vitro* populations in culture.

### 4.5 IBD dataset experiments

We downloaded a dataset of 1304 metagenomic samples from 106 individuals as part of a case-control study investigating irritable bowel disease [3]. We generated log_2_(PTR) estimates using CoPTR. Sequencing reads were mapped to the IGGdb [22] database of representative genomes from high-quality MAGs, assemblies, and complete reference genomes selected from 95% ANI clusters using CoPTR’s wrapper around Bowtie2.

#### Computing the fraction of variance explained

Let *r*_*ij*_ be the *j*-th PTR of a species observed in categorical variable *i* (i.e. an individual, age group, sex, or disease status). To compute the fraction of variance explained we fit the random effects model

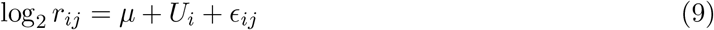

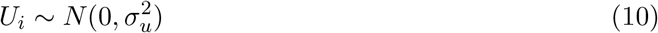

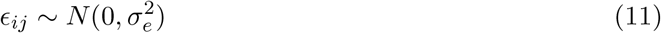

and reported 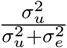 per species. Because individuals accounted for a large fraction of variation, we selected one PTR at random from each individual to estimate variance components for disease status, age, and sex. For age, we divided individuals into a younger and older group using 18 years as a cutoff.

#### Correlation with relative abundances

We computed correlation with relative abundances in two ways. We matched species names from PTRs to estimates from MetaPhlAn2 [24], and computed relative abundances from the read counts mapped using CoPTR. For each species with more than 25 PTRs, we computed standardized log_2_(PTR) and standardized log_2_(Rel Abun) by subtracting the mean and dividing by the standard deviation, then concatenated the resulting estimates from all species together.

#### Associating PTRs with disease status

Because individuals have multiple samples, PTR estimates from the same individual are not independent. Therefore, we tested for a difference in means between cases in controls by taking the mean per individual and adjusting by sample size. We chose this strategy over a linear mixed model because it has higher statistical power. Let *r*_*ij*_ be the *j*-th estimate of a PTR in a species for individual *i*, let *n*_*i*_ be the total number of PTRs in individual *i* for that species, and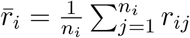 We fit the model

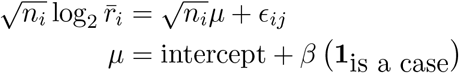

We computed *p*-values separately for each species and disease status, and adjusted for the false discoveries using the Benjamini-Hochberg procedure [34]. We limited our investigation to species with at least 10 PTR estimates in both cases and controls.

#### Associating PTRs with relative abundances and metabolomics

Because relative abundances and metabolite quantities change per sample, we could not use the same association procedure. We therefore used the linear mixed model

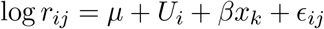

where *µ* is a fixed mean, *U*_*i*_ is a random effect for each individual, *x*_*k*_ is the measurement of interest (a relative abundance or metabolite quantity). For metabolites, we used a log transformation with pseudo count 1 for zeros following the original study [3]. For metabolites, we limited our associations to named metabolites in the Human Metabolome Database. *p*-values were adjusted for false-discoveries using the Benjamini-Hochberg procedure [34].

## Acknowledgements

This material is based upon work supported by the National Science Foundation Graduate Research Fellowship to T.A.J. under Grant No. DGE-1644869. T.K. is a CIFAR Azrieli Global Scholar in the Humans & the Microbiome Program. Additional support was provided by NIH/NCI Grant No. U54CA209997 Driving Biological Projects and Columbia University’s 2020/2021 Data Science Institute Seed Grant.

