## Supplementary Material for "Accurate and robust inference of microbial growth dynamics from metagenomic sequencing"

#### 1 Supplementary Note: PTRs measure DNA replication and generation time

Here we adapt an argument from Bremer and Churchward [1] to show that PTRs provide information about DNA replication and generation time. We need two parameters

1. $C$: The time required to replicate the bacterial chromosome after replication begins.
2. $\tau$: The generation time.

In microbiology, the generation time is defined to be the population doubling time under exponential growth. If the population size at time $t$ is given by

$$N(t) = N_0 2^{t/\tau},$$

the generation time is equivalent to the cell doubling time, and is inversely proportional to the log population growth rate $\frac{\ln 2}{\tau}$.

Most bacteria have a single circular chromosome. Replication begins at a single replication origin, and two replication forks move in either direction along the chromosome until the replication terminus. Under the above model Bremer and Churchward [1] showed that that the ratio of average number of replication origins in a cell $\bar{I}$ to average replication terimini $\bar{T}$

$$\bar{I}/\bar{T} = 2^{C/\tau}.$$

Therefore

$$\log_2\left(\bar{I}/\bar{T}\right) = \frac{C}{\tau}.$$

We want to show this hold in general—regardless of the form of $N(t)$. The argument has three steps. First, we will compute the average rate of DNA synthesis. Second, we will fix the rate of DNA synthesis to compute how many genomes will be added over an interval of time. Third, given the number of genomes added over an interval we can solve for the generation time using its definition.

Let $I(t)$ be the number of replication origins at time $t$, and let $T(t)$ be the number of replication termini. Once DNA replication begins, the two strands of the chromosome separate, and replication forks proceed along both sides of the chromosome. This means $T(t)$ gives the number of complete genomes in the population, since once replication finishes a new completed genome, and therefore terminus, has been added. It follows that the number of replication forks at time $t$ is given by

$$2(I(t) - T(t)).$$

The rate of chromosome replication is essentially constant [2], so each fork produces DNA at a rate of $\sim \frac{1}{2C}$. Therefore, the rate of DNA synthesis in the population is

$$\frac{1}{C}\left(I(t) - T(t)\right).$$

Since $T(t)$ gives the number of complete genomes, dividing by $T(t)$ gives the rate of DNA synthesis per genome

$$\frac{\frac{1}{C}\left(I(t) - T(t)\right)}{T(t)} = \frac{1}{C}\left(R(t) - 1\right)$$
$$R(t) := \frac{I(t)}{T(t)} = \frac{\bar{I}(t)}{\bar{T}(t)}.$$

$R(t)$ corresponds to the PTR. This demonstrates that $R(t)$ is correlated with the average rate of DNA synthesis.

Given a particular time $t_0$ we want to compute how many genomes will be added over an interval. For convenience, let $T_0 = T(t_0)$ and $R_0 = R(t_0)$. The term $\frac{1}{C}(R_0 - 1)$ says that each genome at $t_0$ adds $R_0 - 1$ genomes over $C$ time. Hence, the number of genomes added over $[t_0, t_0 + C]$ is equal to the current number of genomes, $T_0$, plus then number of genomes added, $T_0(R_0 - 1)$. Therefore

$$\#\text{ genomes added over } [t_0, t_0 + C] = T_0 + T_0(R_0 - 1) = T_0 R_0$$

Note that the equation says nothing about genomes removed during this period.

Now we want to apply the definition of generation time. The definition states that generation time is equivalent to population doubling under exponential growth. Thus, we want to know how long it would take for the number of genomes to double, $\tau_0$, given a fixed $R_0$. Treating the number of genomes as continuous, we can write

$$T\left(t_0 + \frac{t}{C}\right) = T_0 R_0^{t/C} \tag{1}$$

So we need to solve

$$2T_0 = T_0 R_0^{\tau_0/C} \implies \log_2 R_0 = \frac{C}{\tau_0}$$

which is what we wanted to show.

A consequence is that under exponential growth

$$T(t) = T_0 R_0^{t/C}.$$

Taking the derivative of $\log_2 T$ we get

$$\frac{d}{dt}\log_2 T(t) = \frac{1}{C}\log_2 R_0.$$

In this specific case, $R(t)$ corresponds to changes in population size.

There are key conceptual differences between our argument and Bremer and Churchward [1]. Bremer and Churchward [1] start with an assumption of the form $N(t)$, and derive expressions for $I(t)$ and $T(t)$ using the parameters $\tau$, $C$, and an additional parameter $D$ that measures the time between genome replication and cell division. Specifically, they assume

$$N(t) = N_0 2^{t/\tau}$$
$$I(t) = I_0 2^{t/\tau}$$
$$T(t) = T_0 2^{t/\tau}$$

Under exponential growth each of these quantities count the same thing but shifted in time, so

$$I_0 = N_0 2^{(C+D)/\tau}$$
$$T_0 = N_0 2^{D/\tau}$$

In contrast, here we do not want to assume a specific form for $N(t)$, $I(t)$ and $T(t)$. In general, we want $I(t)$ and $T(t)$ to arbitrary, and reflect some underlying model of dynamics. We derive an expression for DNA synthesis under arbitrary $I(t)$ and $T(t)$. We solve for $\tau(t)$ by applying the definition of generation time.

### 2 Supplementary Note: Modeling the density of reads along the genome

The results of the previous section demonstrated that $\log_2(\bar{I}/\bar{T}) = \frac{C}{\tau}$, where $\bar{I}$ is the average number of copies of the replication origin in a population, and $\bar{T}$ is the average number of copies of the replication terminus (we have dropped the explicit dependence on $t$ for notation). Suppose we are interested in the ratio of the average copies of an arbitrary position $A$ along the chromosome to the replication terminus. Let $\bar{A}$ be the average copies of $A$, and let $C_A$ be the time it takes the replication fork to move from $A$ to the replication terminus. Note that before the replication fork crosses $A$ there is only one copy, and after the fork crosses $A$ there are two copies. Thus, replacing $I$ with $A$ and $C$ with $C_A$ in the previous argument shows that

$$\frac{1}{C_A}(A - T)$$

also gives the rate of DNA synthesis. Therefore

$$\log_2(\bar{A}/\bar{T}) = \frac{C_A}{\tau}$$

If we assume that chromosome replication happens at a constant rate along the genome, then $C_A$ depends on the distance from $A$ to the replication terminus. Let $b$ be the shortest number of bases between the origin and $A$, such that if we move from origin to the $A$ we do not need to cross the terminus. Let $d$ be the number of bases from the origin to the terminus, and define $f = b/d$. Then $C_A = C(1 - f)$. Rearranging terms from above, we have

$$\log_2(\bar{T}) = \log_2(\bar{I}) - \frac{C}{\tau}$$
$$\log_2(\bar{T}) = \log_2(\bar{A}) - \frac{C_A}{\tau} = \log_2(\bar{A}) - \frac{C(1-f)}{\tau}$$

Subtracting the first equation from the second, and rearranging terms

$$\implies \log_2(\bar{A}) = \log_2(\bar{I}) - \frac{Cf}{\tau}$$

Consequentially, the average copies of position $A$ decays log-linearly with distance from the replication origin.

This also means that any probabilistic model of reads along the genome, coverage should decay log-linearly away from the replication origin. Therefore, we propose the following model. Let $[0, 1]$ represent coordinates along a continuous approximation of a reference genome. Thus 0 is the beginning of the reference, and 1 is the end. The model parameters are the origin position $x_i$, terminus position $x_t = (x_i + 0.5) \bmod 1$, and PTR $r$. We want a probability density given by

$$\alpha = \frac{\log_2 r}{x_i - x_t} = \frac{\log_2 p(x_i) - \log_2 p(x_t)}{x_i - x_t}$$

$$x_1 = \min\{x_i, x_t\}$$

$$x_2 = \max\{x_i, x_t\}$$

$$c(x) = \begin{cases} \log_2 p(x_i) \text{ if } x = x_i \\ \log_2 p(x_t) \text{ if } x = x_t \end{cases}$$

$$\log_2 p(x) = \begin{cases} -\alpha(x - x_1) + c(x_1) & \text{if } x \leq x_1 \\ \alpha(x - x_1) + c(x_1) & \text{if } x_1 < x < x_2 \\ -\alpha(x - x_2) + c(x_2) & \text{if } x \geq x_2 \end{cases}$$

We need to compute $\log p(x_i)$ and $\log p(x_t)$ such that

$$\int_0^1 2^{\log_2 p(x)} = 1.$$

We can use the integral, and the constraint that $\log_2 p(x_i) - \log_2 p(x_t) = \log_2 r$ to solve for each. There are two cases. If $x_i \leq x_t$, then

$$\log_2 p(x_i) = \log_2 \ln 2 - \log_2 \left( \frac{1}{\alpha} \left[ 2^{\alpha x_1} + 2^{\alpha(x_2 - x_1)} - 2^1 - 2^{-\alpha(1-x_2) - \log_2 r} + 2^{-\log_2 r} \right] \right)$$

$$\log_2 p(x_t) = \log_2 p(x_i) - \log_2 r$$

If $x_t < x_i$, then

$$\log_2 p(x_t) = \log_2 \ln 2 - \log_2 \left( \frac{1}{\alpha} \left[ 2^{\alpha x_1} + 2^{\alpha(x_2 - x_1)} - 2^1 - 2^{-\alpha(1-x_2) + \log_2 r} + 2^{+\log_2 r} \right] \right)$$

$$\log p_2(x_i) = \log_2 p(x_t) + \log_2 r$$

### 3 Supplementary Note: Variational inference for multi-mapped reads

Suppose we have the following model for drawing the assignment of sequencing reads from a set of $\mathcal{G}$ reference genomes indexed from $1...g$.

1. Draw probabilities that a read originates from a reference genome:

$$\pi \sim \text{Dirichlet}(\alpha_1, ...., \alpha_g).$$

2. For each read $i = 1...n$, pick a reference genome:

$$z_i|\pi \sim \text{Categorical}(\pi)$$

For notation, let $z_i$ be an indicator vector, where $z_{ij} = 1$ if $z_i$ is assigned to genome $j$. Let $x_i = (x_{i1}, x_{i2}, ..., x_{ig}) \in \{0, 1\}^g$ where $x_{ij} = 1$ if the read maps to a position in genome $j$, and is 0 otherwise. If read $i$ maps to only one genome, then $z_i = x_i$. If read $i$ maps to multiple genomes, then $z_i$ places a restriction on $x_i$: if $z_{ij} = 1$ then it must be true that $x_{ij} = 1$—assuming that one of the given mappings is always correct. Thus, we can model $x_{ij}$ as

$$p(x_{ij} = 1|z_{ij}) = \begin{cases} 1 & \text{if } z_{ij} = 1 \\ \rho_{ij} & \text{otherwise.} \end{cases}$$

Now consider that once a sequencing read is observed, all valid mappings are determined. Hence $\rho_{ij} = 1$ if read $i$ has a valid mapping to genome $j$ and 0 otherwise. Therefore

$$p(x_i|z_i) = \prod_{j=1}^{g} x_{ij}^{z_{ij}}$$

if we define $0^0 = 1$.

If we could compute

$$p(z_{1:n}, \pi|x_{1:n})$$

then we could assign reads to genomes based on the posterior. However, the normalizing constant is

$$p(x_{1:n}) = \int_{\pi} \sum_{z_{1:n}} p(x_{1:n}, z_{1:n}, \pi)\, d\pi$$

which requires summing over an exponential number of combinations of $z_{1:n}$. Nonetheless, we can compute

$$p(z_i|\pi, x_i) = \frac{\prod_{j:x_{ij}=1} \pi_j^{z_{ij}}}{\sum_{j:x_{ij}=1} \pi_j}$$

Additionally, since the Dirichlet distribution is a conjugate prior for the multinomial distribution

$$p(\pi|z_{1:n}, x_{1:n}) = p(\pi|z_{1:n}) = \text{Dirichlet}\left(\pi; \alpha + \sum_{i=1}^{n} z_i\right)$$

Therefore we can compute all of the complete conditionals. This means we can approximate $p(z_{1:n}, \pi|x_{1:n})$ either using Gibbs sampling or mean field variational inference [3]. We chose variational inference.

Variational inference approximates an intractable posterior one by a tractable one $q$ whose parameters are optimized to minimize the a lower bound on the log-likelihood. Equivalently, variational inference minimizes Kullback-Leibler divergence between the true posterior and the approximation. For the mean field approximation, $q(z_{1:n}, \pi) = q(\pi) \prod_{i=1}^{n} q(z_i)$. The optimal choice for an approximation $q(z_i)$ is given by (see Blei et al. [3])

$$\begin{aligned} q(z_i) &\propto \exp\left\{\mathbb{E}_{-z_i}\left[\log p(z_i|\pi, x_i)\right]\right\} \\ &\propto \exp\left\{\sum_{j:x_{ij}=1} z_{ij}\, \mathbb{E}_{-z_i}[\log \pi_j]\right\} \end{aligned}$$

This set of equations give the natural parameters of a multinomial distribution, so $q(z_i) = \text{Multinomal}(z_i; 1, \phi_i)$ where $\log \phi_{ij} = \mathbb{E}_{-z_i}[\log \pi_j] + \textit{const}$ if $x_{ij} = 1$, and $\phi_{ij} = 0$ otherwise. The optimal choice for $q(\pi)$ is given by

$$\begin{aligned} q(\pi) &\propto \exp\left\{\mathbb{E}_{-\pi}\left[\log p(\pi|z_{1:n})\right]\right\} \\ &\propto \exp\left\{\mathbb{E}_{-\pi}\left[\sum_{i=1}^{n} \log p(z_i|\pi)\right] + \log p(\pi)\right\} \\ &\propto \exp\left\{\sum_{i=1}^{n} \sum_{j:x_{ij}=1} \phi_j \log \pi_j + \sum_{j=1}^{g} (\alpha_j - 1) \log \pi_j\right\} \\ &\propto \exp\left\{\sum_{i=1}^{n} \sum_{j=1}^{g} \phi_j \log \pi_j + \sum_{j=1}^{g} (\alpha_j - 1) \log \pi_j\right\} \\ &= \text{Dirichlet}\left(\pi; \alpha + \sum_{i=1}^{n} \phi_i\right) \end{aligned}$$

The second set of equations gives the natural parameters of a Dirichlet distribution, so $q(\pi) = \text{Dirichlet}(\pi; \eta)$. Now we can compute for the final expectation:

$$\mathbb{E}_{-z_i}[\log \pi_j] = \Psi(\eta_j) - \Psi\left(\sum_{j'=1}^{g} \eta_{j'}\right).$$

The final component is computing the variational objective function. This is given by

$$L(z_{1:n}, x_{1:n}, \pi; \phi_{i:n}, \eta) = \mathbb{E}_q[\log p(z_{1:n}, x_{1:n}, \pi)] - \sum_{i=1}^{n} \mathbb{E}_q[\log q(z_i; \phi_i)] - \mathbb{E}_q[\log q(\pi; \eta)]$$

The second two terms are the entropy of a multinomial and Dirichlet distribution respectively. The joint model likelihood is

$$p(z_{1:n}, x_{1:n}, \pi) = p(\pi) \prod_{i=1}^{n} \prod_{j=1}^{g} x_{ij}^{z_{ij}} \pi_j^{z_{ij}}$$

Hence

$$\mathbb{E}_q[\log p(z_{1:n}, x_{1:n}, \pi)] = \mathbb{E}_q[\log p(\pi)] + \sum_{i=1}^{n} \sum_{j=1}^{g} \mathbb{E}_q[z_{ij}] x_{ij} + \mathbb{E}_q[z_{ij}]\, \mathbb{E}_q[\log \pi_j]$$

We now can define an inference procedure for computing the approximate posterior.

1. Initialize variational parameters $\phi_{1:n}$ and $\eta$.
2. While $L(z_{1:n}, x_{1:n}, \pi; \phi_{i:n}, \eta)$ has not converged:
   (a) Set $q(z_i) \propto \exp\left\{\sum_{j:x_{ij}=1} z_{ij}\, \mathbb{E}_{-z_i}[\log \pi_j]\right\}$ for $i = 1...n$
   (b) Set $q(\pi) = \text{Dirichlet}\left(\pi; \alpha + \sum_{i=1}^{n} \phi_i\right)$

The $q(z_i)$ define an approximate posterior over $z_i$.

### 4 Supplementary Figures

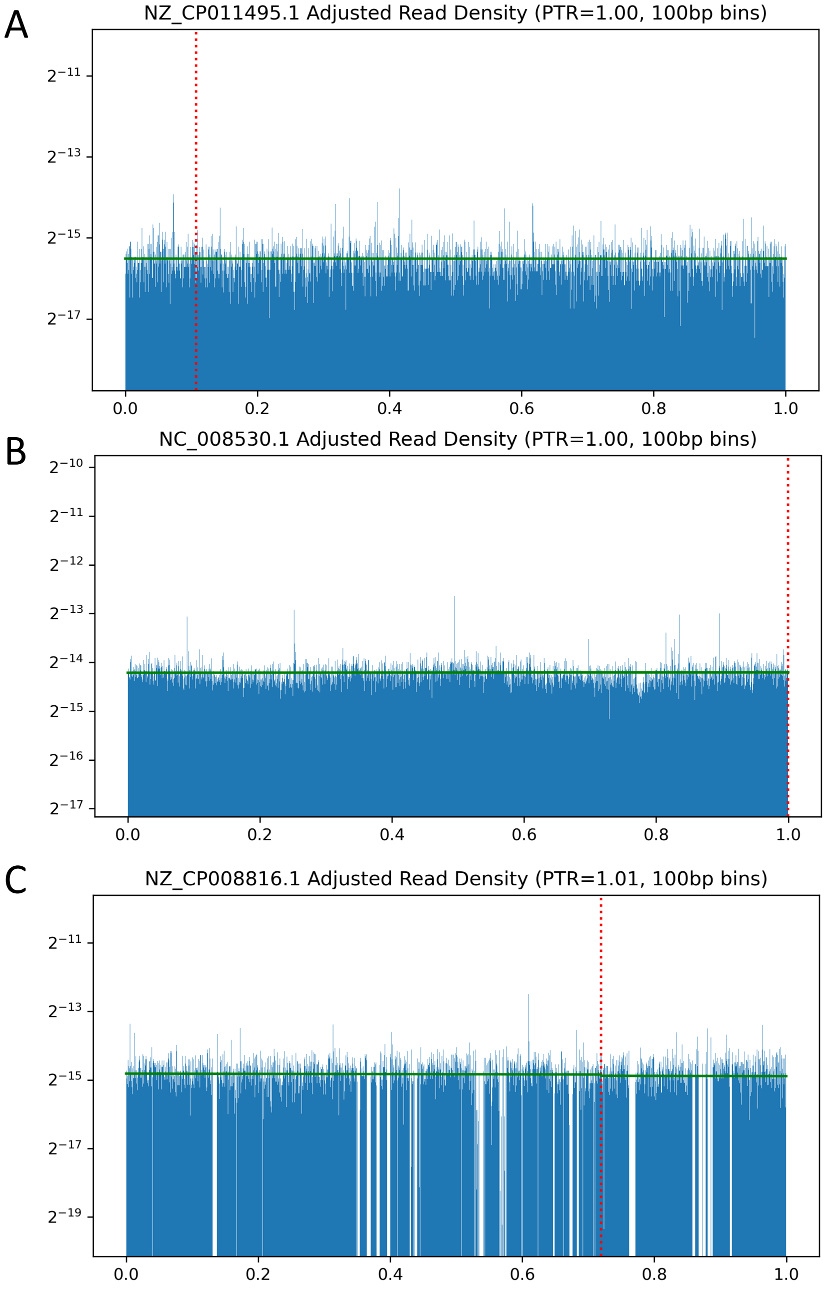


**Supplementary Figure 1: Adjusted coverage maps used for simulated data.** Coverage maps in 100bp windows for three species: **(A)** E. coli, **(B)** L. gasseri, and **(C)** E. faecalis. Reported PTRs are computed after adjustment for the estimated PTR on each dataset. Dashed red lines denote the replication origin from the DoriC database [4].

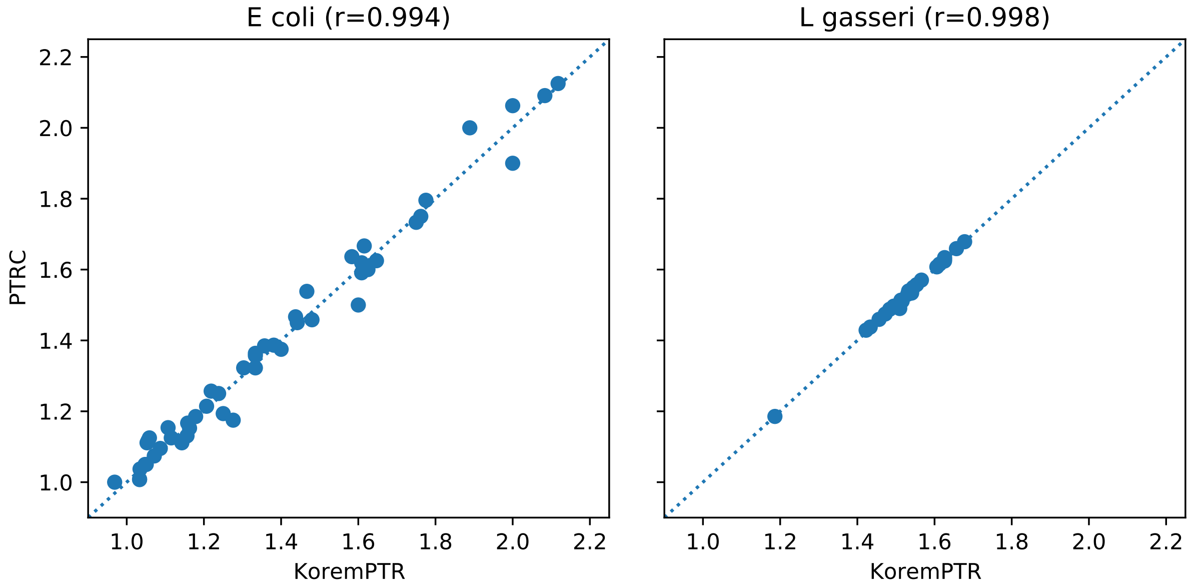


**Supplementary Figure 2: Comparison between PTRC and KoremPTR on two datasets.** Left: 55 genomic samples of *E. coli*. Right: 25 genomic samples of *L. gasseri*.

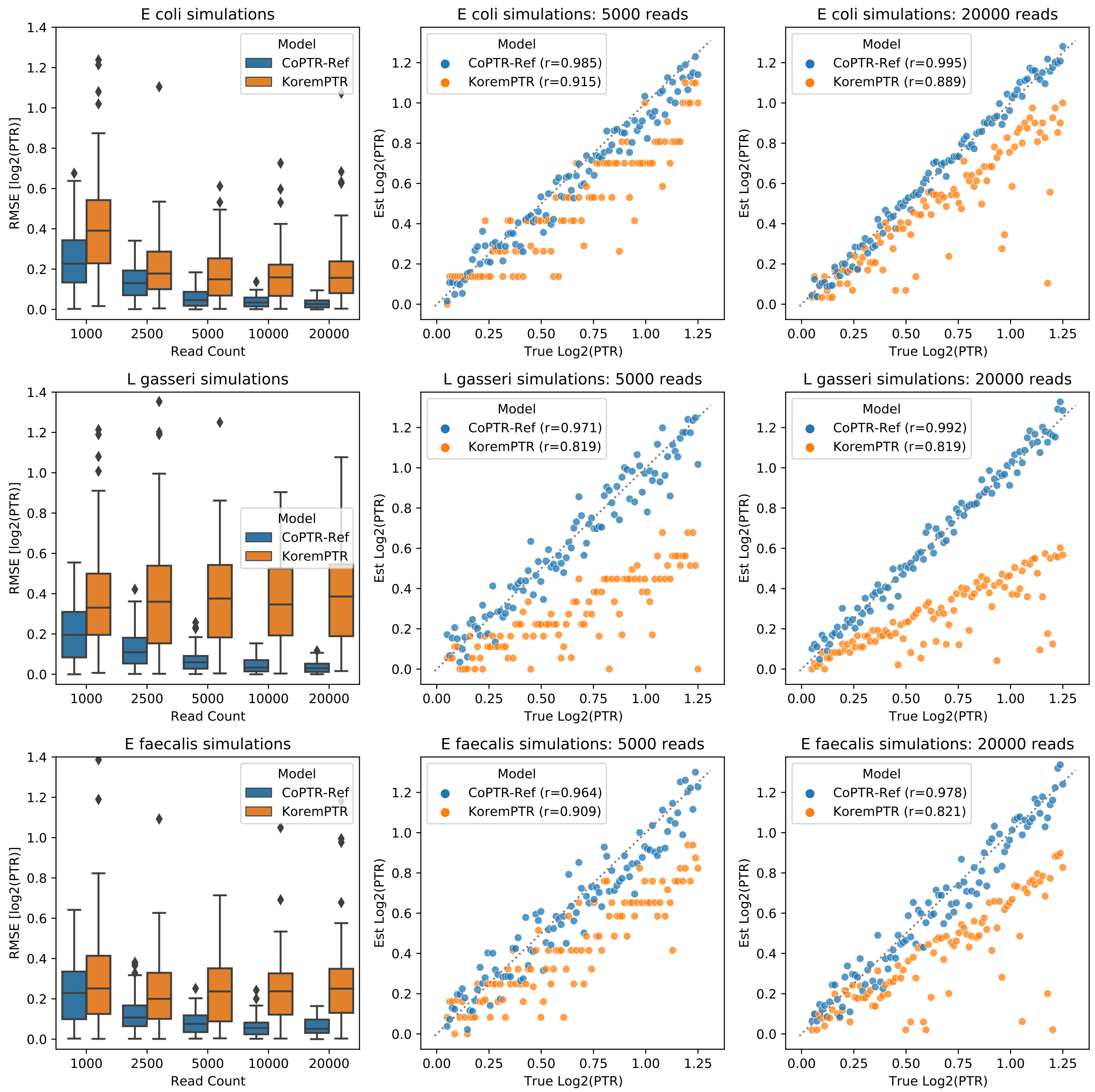


**Supplementary Figure 3: Simulation comparison between CoPTR-Ref and KoremPTR**. Top: Simulations based on the *E. coli* density map. The first and last columns recapitulate Figure 2. Middle: Simulations based on the *L. gasseri* density map. Bottom: Simulations based on the *E. faecalis* density map.

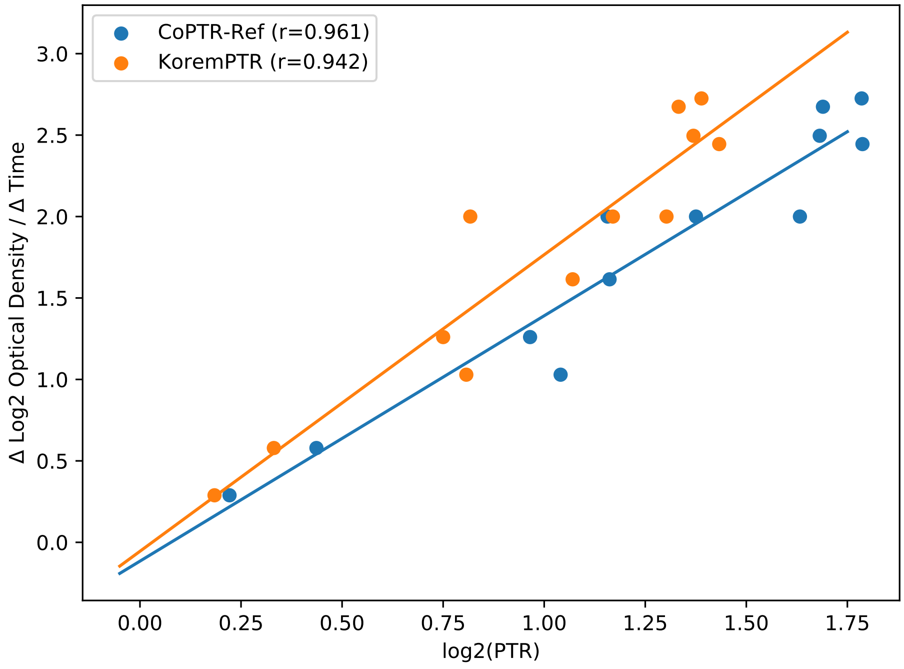


**Supplementary Figure 4: Correlation between log PTR and changes in log abundances for *E. coli* grown in culture.** Abundance is measured using the optical density of the culture.

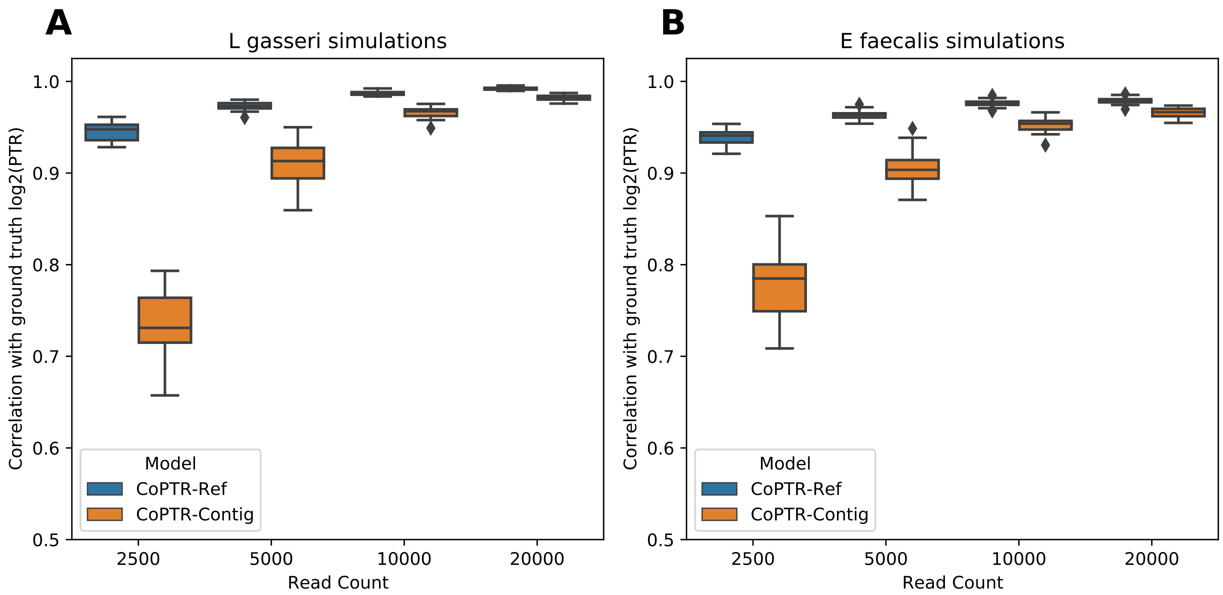


**Supplementary Figure 5: Simulation comparison between CoPTR-Ref and CoPTR-Contig**. **(A)** Comaparison on simulations from *L. gasseri* density maps. **(B)** Comparison on simulations from *E. faecalis* density maps.

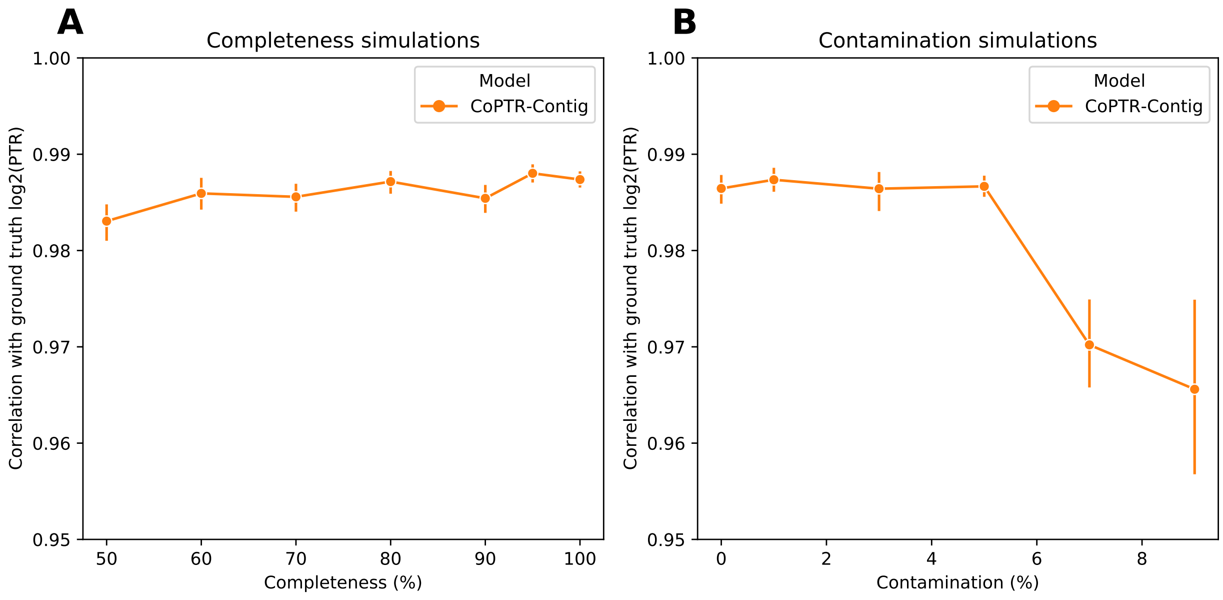


**Supplementary Figure 6: CoPTR-Contig is robust to genome completeness and contamination.** Evaluation of genome completeness **(A)** and contamination **(B)** on PTR estimation. Error bars depict one standard deviation across 20 simulation replicates, each replicate consisting of 100 simulated PTRs from a simulated assembly.

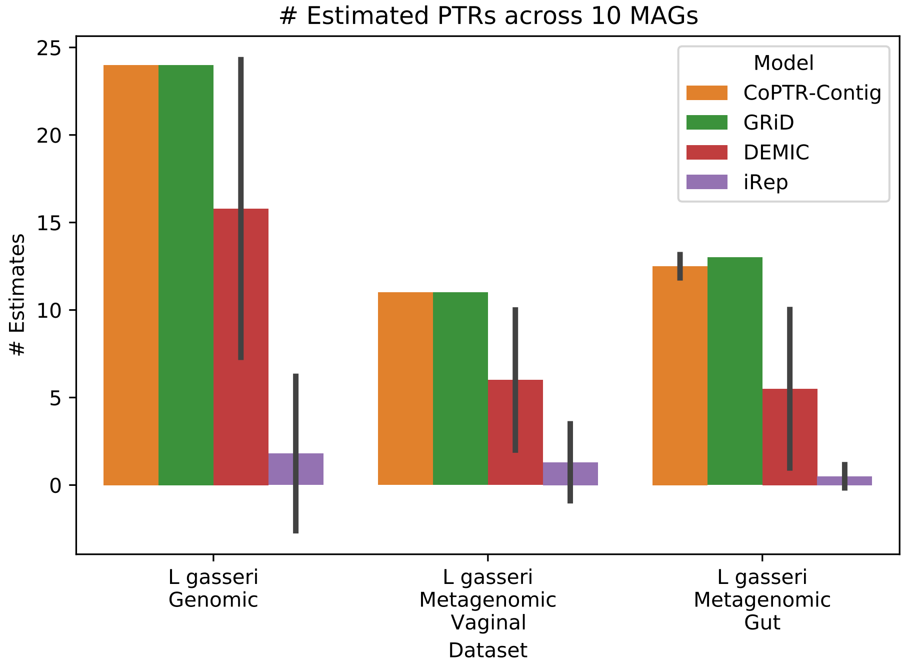


**Supplementary Figure 7: Number of PTR estimates output by each model.** Mean number of PTR estimates across 10 high quality MAGs. Error bars depict one standard deviation.

### 5 Supplementary Tables

| Parameter | Effect Size | $p$-value |
|---|---|---|
| Erythromycin | $-0.3412$ | $1.40 \times 10^{-04}$ |
| Nalidixic acid | $0.2819$ | $3.99 \times 10^{-04}$ |
| $a_{treatment}$ | $-0.0629$ | $1.67 \times 10^{-03}$ |
| $b_{treatment}$ | $1.5183$ | $8.08 \times 10^{-34}$ |
| $a_{recovery}$ | $-0.1939$ | $9.38 \times 10^{-02}$ |
| $b_{recovery}$ | $1.1945$ | $2.51 \times 10^{-24}$ |
| $a_{stationary}$ | $-0.0902$ | $1.46 \times 10^{-01}$ |
| $b_{stationary}$ | $0.6591$ | $1.78 \times 10^{-13}$ |

**Supplementary Table 1: Effect sizes and p-values from the antibiotic resistance experiment.**

| Species | Effect Size | $p$-value | $q$-value | # Cases | # Controls |
|---|---|---|---|---|---|
| Subdoligranulum sp. | -0.157350 | 0.000210 | 0.025177 | 29 | 20 |
| COE1 sp. | 0.114031 | 0.001040 | 0.062412 | 11 | 12 |
| Dorea longicatena | 0.176232 | 0.003077 | 0.123098 | 58 | 25 |
| Flavonifractor plautii | 0.050461 | 0.007306 | 0.202415 | 47 | 22 |
| ER4 sp. | -0.051200 | 0.009973 | 0.202415 | 14 | 15 |

**Supplementary Table 2: Top 5 most significantly associated PTRs with Crohn's disease.** Total of 120 hypothesis tests were performed. Number of cases and controls refer to individuals, not total samples.

| Species | Effect Size | $p$-value | $q$-value | # Cases | # Controls |
|---|---|---|---|---|---|
| Roseburia intestinalis | 0.094214 | 0.000010 | 0.001068 | 27 | 23 |
| Ruminiclostridium sp. | -0.138431 | 0.000505 | 0.025750 | 12 | 18 |
| Subdoligranulum sp. | -0.168006 | 0.000792 | 0.026915 | 19 | 20 |
| Faecalibacterium HGM13285 | 0.742445 | 0.003055 | 0.077897 | 27 | 24 |
| Flavonifractor plautii | 0.052042 | 0.012679 | 0.258653 | 27 | 22 |

**Supplementary Table 3: Top 5 most significantly associated PTRs with Ulcerative Colitis.** Total of 102 hypothesis tests were performed. Number of cases and controls refer to individuals, not total samples.

| Species | Effect Size | $p$-value | $q$-value | Sample Size |
|---|---|---|---|---|
| Oscillibacter sp KLE 1745 | -1.632241 | 0.000003 | 0.000342 | 45 |
| Actinobacillus unclassified | 63.891339 | 0.000262 | 0.016633 | 57 |
| Peptostreptococcaceae noname unclassified | 10.137852 | 0.000908 | 0.038427 | 288 |
| Haemophilus parainfluenzae | 0.861744 | 0.010901 | 0.298997 | 386 |
| Alistipes onderdonkii | -0.628384 | 0.011772 | 0.298997 | 331 |

**Supplementary Table 4: Top 5 most significantly associated relative abundances with *R. intestinalis*.** Total of 127 hypothesis tests were performed.

| Metabolite | HMDB Accession | Effect Size | $p$-value | $q$-value | Sample Size |
|---|---|---|---|---|---|
| 2-hydroxyglutarate | HMDB59655 | 0.028988 | 0.000194 | 0.042292 | 185 |
| 13-cis-retinoic acid | HMDB06219 | 0.010219 | 0.000195 | 0.042292 | 131 |
| malonate | HMDB00691 | 0.008374 | 0.001990 | 0.280461 | 166 |
| C36:0 DAG | HMDB07158 | -0.009536 | 0.002591 | 0.280461 | 118 |
| C20:4 LPE | HMDB11517 | 0.033313 | 0.003635 | 0.282199 | 185 |

**Supplementary Table 5: Top 5 most significantly associated metabolites with *R. intestinalis*.** Total of 433 hypothesis tests were performed.

| Description | Download Source | Identifiers |
|---|---|---|
| *E. coli* genomes | NCBI RefSeq | NC_002695.2, NC_004431.1, NC_007779.1, NC_010468.1, NC_010498.1 |
| *L. gasseri* genomes | NCBI RefSeq | NC_008530.1, NZ_CP006803.1, NZ_CP021427.1, NZ_CP054875.1 |
| *E. faecalis* genomes | NCBI RefSeq | NC_004668.1, NC_018221.1, NZ_CP008816.1, NZ_CP018004.1, NZ_CP018102.1 |
| *E. coli* MAGs | IGGdb | ERS235517_65, ERS235554_22, ERS235558_17, ERS235567_48, ERS235580_5, ERS235591_41, ERS235592_93, ERS235593_34, ERS235594_3, ERS235598_31 |
| *L. gasseri* MAGs | IGGdb | ERS396471_42, ERS473039_24, ERS473053_7, ERS473384_2, ERS608476_13, ERS608506_43, SRS1719133_5, SRS1735437_2, SRS255102_17, SRS294994_4 |
| *E. coli* genomic dataset | NCBI SRA | ERR969279, ERR969280, ERR969281, ERR969282, ERR969283, ERR969284, ERR969285, ERR969286, ERR969287, ERR969288, ERR969289, ERR969290, ERR969291, ERR969292, ERR969293, ERR969294, ERR969295, ERR969296, ERR969297, ERR969298, ERR969299, ERR969300, ERR969301, ERR969302, ERR969303, ERR969304, ERR969305, ERR969306, ERR969307, ERR969308, ERR969309, ERR969310, ERR969311, ERR969312, ERR969313, ERR969314, ERR969315, ERR969316, ERR969317, ERR969318, ERR969319, ERR969320, ERR969321, ERR969322, ERR969323, ERR969324, ERR969326, ERR969327, ERR969328, ERR969329, ERR969330, ERR969331, ERR969332, ERR969333, ERR969334 |
| *E. coli* metagenomic dataset | NCBI SRA | SRR1952457, SRR1952459, SRR1952575, SRR1952582, SRR1952598, SRR1952613, SRR1952621, SRR2077403, SRR2175645, SRR2175647, SRR2175654, SRR2175658, SRR2175659, SRR2175725, SRR2175726, SRR2175755, SRR2175766, SRR2175767, SRR2175777, SRR2175780, SRR2175792, SRR2240290, SRR2240728, SRR2241023, SRR2241109, SRR2241118 |
| *L. gasseri* genomic dataset | NCBI SRA | ERR969426, ERR969427, ERR969428, ERR969429, ERR969430, ERR969431, ERR969432, ERR969433, ERR969434, ERR969435, ERR969436, ERR969437, ERR969438, ERR969440, ERR969441, ERR969442, ERR969443, ERR969444, ERR969445, ERR969446, ERR969447, ERR969448, ERR969449, ERR969450 |
| *L. gasseri* vaginal metagenomic dataset | NCBI SRA | SRR059392, SRR059393, SRR059473, SRR063749, SRR1804974, SRR2175824, SRR2241374, SRR513144, SRR513147, SRR514852, SRR628270 |
| *L. gasseri* gut metagenomic dataset | NCBI SRA | SRR5935920, SRR5936055, SRR5936136, SRR5936140, SRR5946677, SRR5946785, SRR5946864, SRR5946962, SRR5947082, SRR5950499, SRR5950538, SRR5950556, SRR5950631 |
| IBD Dataset | NCBI SRA | Available under BioProject PRJNA398089 |

**Supplementary Table 6: Accession numbers and identifiers of genomes and datasets used.**

[4] Hao Luo and Feng Gao. Doric 10.0: an updated database of replication origins in prokaryotic genomes including chromosomes and plasmids. *Nucleic acids research*, 47(D1):D74–D77, 2019.
